# An integrative proteotranscriptomics approach reveals new ADAM9 substrates and downstream pathways

**DOI:** 10.1101/2024.10.01.616047

**Authors:** Congyu Lu, Xiaolu Xu, Neha Sindhu, Jessica Rainey, Yuhan Zhang, Shawn W. Polson, Jing Qiu, Shuo Wei

## Abstract

The disintegrin metalloprotease ADAM9 is a cell-surface protease that can shed the ectodomain of membrane protein substrates. Dysregulated ADAM9 activity has been implicated in several diseases such as solid tumors, autoimmunity, inflammatory diseases, and COVID-19. Despite its importance, the substrates and targets of ADAM9 in normal and pathological processes are poorly understood. Here, we developed an integrative proteotranscriptomics approach to systematically identify the transcriptional and post-transcriptional targets of ADAM9 in HCT116 cells, which have a stable diploid karyotype suitable for omics analyses. Using this approach, we uncovered major signaling pathways downstream of ADAM9, including the oncogenic mTOR pathway and the tumor suppressor FOXO pathway. We also identified several direct and indirect substrates for ADAM9, which may mediate the pathophysiological roles of this protease. This study provides new mechanistic insights into the function of ADAM9 as well as a method that can be applied to other membrane proteases.

## Introduction

Members of the disintegrin metalloproteinase (ADAM) family are key regulators of cellular events such as signaling, adhesion and migration. About half of the ADAMs are proteolytically active and primarily function as enzymes that catalyze ectodomain shedding of membrane proteins (1, 2). Among these, the canonical “sheddase” ADAM9 has emerged as a potential therapeutic target for several diseases. ADAM9 is highly expressed in multiple types of solid tumors and promotes tumor progression (3). Increased ADAM9 expression was also found in chronic obstructive pulmonary disease (COPD) patient samples, and knockout (KO) of *Adam9* protects mice from developing COPD (4). In addition, patients with autoimmune diseases such as systemic lupus erythematosus have higher levels of ADAM9 in their CD4^+^ T cells, and KO of *Adam9* in mice mitigates autoimmunity by reducing the ability of CD4^+^ T cells to differentiate into inflammation-promoting Th17 cells (5). Recently, ADAM9 was reported to be upregulated in young patients with critical COVID-19 and facilitate viral infection and replication (6, 7). Conversely, loss of ADAM9 can lead to retinal degeneration in humans and other mammals (8, 9). However, despite the important roles of ADAM9 in the various diseases, the substrates and targets of this versatile protease remain largely unknown (10). This information is not only pivotal for understanding the mechanisms of action for ADAM9 in these diseases, but also for assessing the potential adverse effects when targeting this sheddase for therapeutic purposes.

Currently, proteomics is essentially the only method for systematic identification of sheddase substrates. Most of the published proteomics studies were focused on the secretome, as the shed ectodomain of substrates can often be detected in the conditioned media of cultured cells or tissues, and their abundance is altered when sheddase activity is manipulated (11). These secretomics-based methods have identified numerous substrates for the “housekeeping” sheddases such as ADAM10 and 17 (2), but do not seem to be effective for other sheddases that may have fewer substrates. For example, recent attempts using secretomics failed to identify any substrates for ADAM15 (12, 13), a sheddase that is similar to ADAM9 in expression and function (14). A possible explanation for the false negative results of these methods is that the shed ectodomain of some substrates may remain attached to the membrane or are quickly turned over, and are hence not present in significant amount in the secretome. One example is the Notch receptors, which are associated with Notch ligands after cleavage by ADAM10, until being internalized and degraded (15). False positive is another problem, as many differentially expressed proteins (DEPs) in the secretome may be regulated through other mechanisms such as transcription, but cannot be distinguished from *bona fide* shedding substrates by secretomics alone. Finally, secretomics provides little information on downstream genes and pathways, which are also important for mediating sheddase function. Therefore, it is necessary to develop new methods to systematically identify sheddase substrates and targets.

The current study aims to establish a new integrative proteotranscriptomics approach to systematically identify the substrates and targets for ADAM9. We carried out RNA-sequencing (RNA-seq) and quantitative proteomics for HCT116 colon cancer cells with and without ADAM9 knockdown (KD; Fig. 1A). HCT116 cells were chosen for this study due to their stable diploid karyotype that is suitable for omics analyses (16). The two omics datasets were integrated using two-way Orthogonal Partial Least Squares (O2PLS) analyses to identify transcriptionally and post-transcriptionally regulated protein targets of ADAM9. Among the post-transcriptional targets that negatively correlate with ADAM9, cell-surface proteins were selected as candidate substrates (Fig. 1B). Using this approach, we successfully uncovered two downstream signaling pathways and three substrates for ADAM9, which may contribute to the pathological roles of this sheddase.

**Figure 1.**
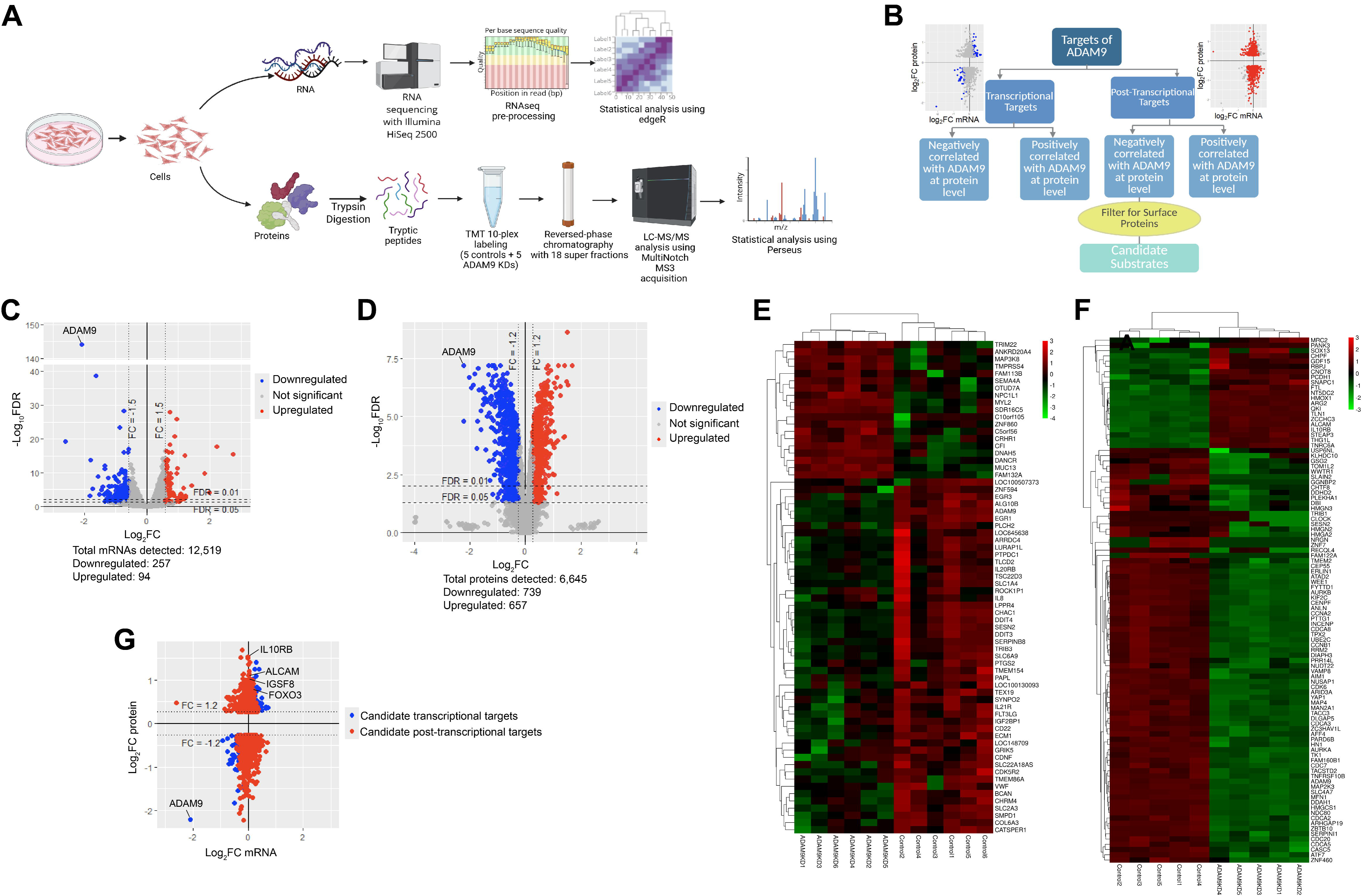
The transcriptome and cellular proteome of HCT116 cells with and without ADAM9 KD. (**A**) Flow chart of our transcriptome and cellular proteome analyses. (**B**) Integration of transcriptome and cellular proteome to identify transcriptional and post-transcriptional targets as well as candidate substrates for ADAM9. (**C, D**) Volcano plots of transcriptome (**C**) and cellular proteome (**D**) showing DEGs (|FC|>1.5, FDR<0.05) and DEPs (|FC|>1.2, FDR<0.05), respectively. ADAM9 is highlighted as a positive control. (**E, F**) Heat maps showing top DEGs (**E**) and DEPs (**F**). Color codes represent z-scores. (**G**) A scatter plot showing the correlation between mRNA and protein levels. |FC|>1.2 at protein level was applied to the identification of candidate transcriptional and post-transcriptional targets to reduce false positives. FC was defined as ADAM9 KD/control.

## Results

### Characterizing and integrating the transcriptome and cellular proteome regulated by ADAM9 in HCT116 cells

We recently showed that ADAM9 expression is elevated in colorectal cancer (CRC) tissues as compared with the adjacent normal tissues taken from the same patients. In HCT116 and SW620 CRC cells, ADAM9 cleaves ephrin-B1 and -B2 to promote downstream Akt kinase activity, thereby facilitating cell migration and invasion (17). To systematically identify other ADAM9 substrates and targets, we knocked down ADAM9 in HCT116 cells using the well characterized siRNA AD9-1 (17). RNA-seq and multiplexed quantitative proteomics with tandem mass tag (TMT) labeling were performed for control and ADAM9 KD cells, and a total of 12,519 mRNAs and 6,645 proteins were detected (Table S1 and S2). Principle component analyses for both omics data show clear segregation of ADAM9 KD samples from the controls (Fig. S1 and S2), and a consistent downregulation of both ADAM9 mRNA and protein was observed (Fig. S3A and B), confirming data quality. Volcano plots of differentially regulated genes (DEGs) and DEPs (FDR<0.05) are shown in Fig. 1C and D, and heat maps of the top DEGs and DEPs in Fig. 1E and F. The transcriptome and proteome data were integrated (Fig. 1G) and the R package OmicsPLS was used to identify the transcriptional and post-transcriptional targets as described in Fig. 1B and Methods. Transcriptional and post-transcriptional targets that positively and negatively correlated with ADAM9 (|FC|>1.2 at protein level) are listed in Table S3 and S4, respectively.

### mTOR and FOXO signaling pathways are downstream of ADAM9 in HCT116 cells

To search for major signaling pathways downstream of ADAM9, we carried out KEGG pathway enrichment analysis for the DEGs (|FC|>1.5), as altered cell signaling often leads to significant changes in the transcriptome. Among the top 10 pathways, 5 are related to Akt (Fig. 2A). Besides the PI3K-Akt pathway, mTOR and FOXO are two major pathways downstream of Akt, and the p53 and MAPK pathways are cross-regulated by Akt (18, 19). A comparison of our DEGs (|FC|>1.5, q<0.05) with those obtained previously using the mTOR inhibitor Torin-1 (20) shows significant overlap (p=1.5E-04; Fig. 2B and Table S5). The validation of mTOR signaling as a target pathway of ADAM9 in HCT116 cells has been published elsewhere (17). Akt also downregulates FOXO signaling by phosphorylating FOXO transcription factors to facilitate their nuclear export and degradation (21). Interestingly, FOXO3, the only FOXO family member detected in HCT116 cells, was one of the top post-transcriptionally upregulated proteins upon ADAM9 KD (FC=1.74, q=2.8E-06). There is also a significant overlap between our DEGs and the genes previously found to be FOXO3 targets (22)(p=0.006; Fig. 2C and Table S6). We confirmed the upregulation of FOXO3 with western blotting using both AD9-1 and a second siRNA targeting ADAM9 (AD9-2; Fig. 2D and S4); no significant alteration of the *FOXO3* mRNA was found (Fig. 2E). Three FOXO3 target genes (*MUC13*, *THBS1*, and *GDA*) were selected from Table S6, and their upregulation following ADAM9 KD was validated using RT-qPCR (Fig. 2E). Consistent with these findings, KD of ADAM9 leads to increased levels of nuclear FOXO3, as shown by immunocytochemistry (ICC; Fig. 2F). Together, these data indicate that ADAM9 suppresses FOXO signaling in HCT116 cells.

**Figure 2.**
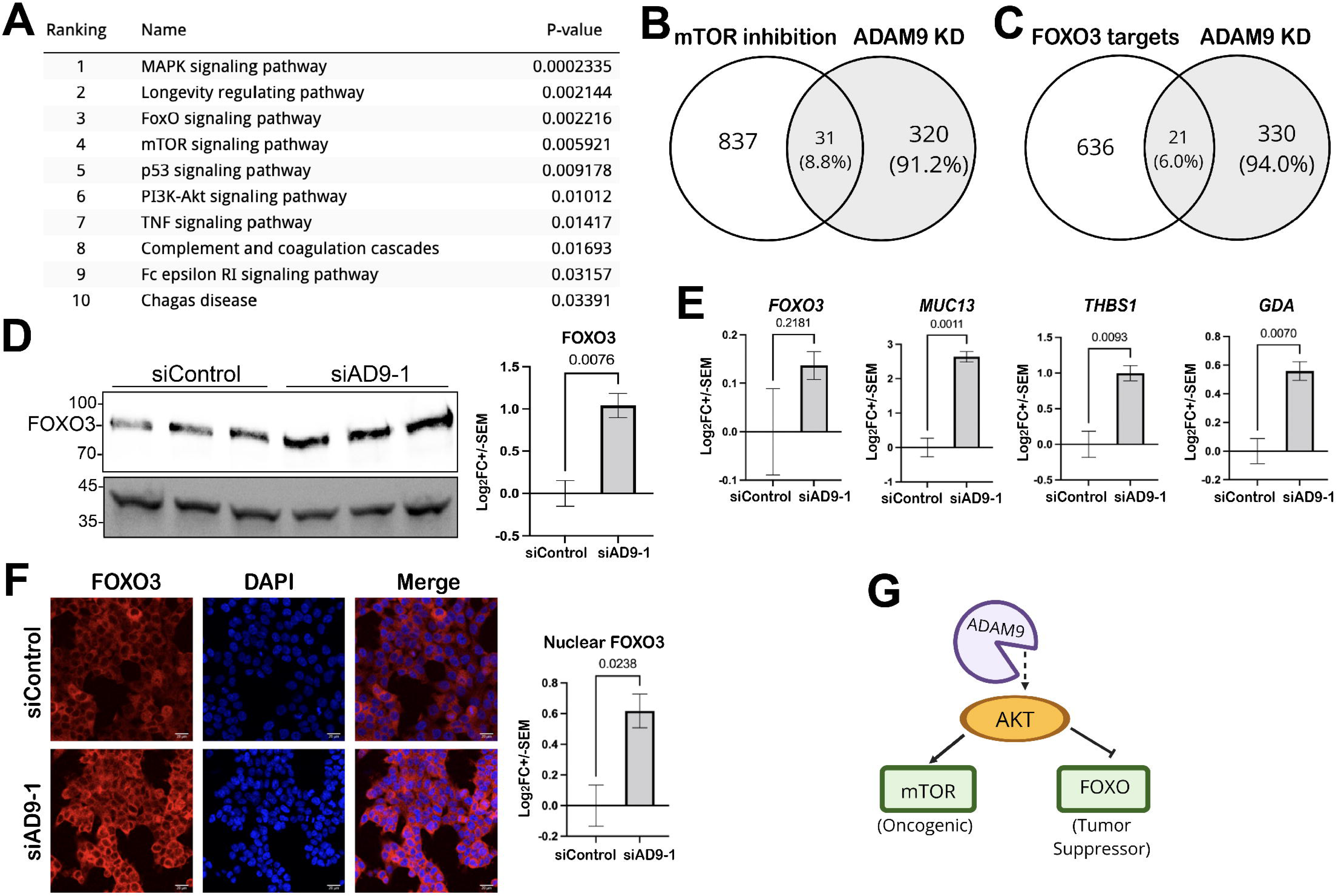
ADAM9 regulates mTOR and FOXO signaling in HCT116 cells. (**A**) KEGG pathway enrichment analysis of transcriptome data showing top 10 pathways affected by ADAM9 KD. (**B**, **C**) Venn diagrams showing significant overlaps between the DEGs found in our transcriptome data and published transcriptional targets of mTOR (**B**) and FOXO3 (**C**). (**D**) Western blots of FOXO3 in control and ADAM9 KD cells (3 biological replicates each). Statistics calculated with unpaired two-tail t test is shown on the right with p-value at the top. (**E**) RT-qPCR results of the indicated genes with and without ADAM9 KD (3 biological replicates each). (**F**) ICC of FOXO3 in control and ADAM9 KD cells. Confocal images were processed with ImageJ to measure the intensity of red fluorescence (FOXO3) within the DAPI-stained area (nucleus). Images of one large field (0.83×0.83 μm) randomly selected from each of the three biological replicates were used for quantification, and unpaired two-tail t test was performed to assess the differences. (**G**) Signaling pathways downstream of ADAM9 identified in HCT116 cells.

### Identifying candidate substrates for ADAM9 from post-transcriptional targets

Upon ADAM9 KD, substrates are expected to accumulate on the membrane unless there are additional feedbacks to regulate their protein levels. Among the post-transcriptional targets that negatively correlate with ADAM9 (|FC|>1.2 for DEP), 188 were annotated as membrane proteins in UniProt. Further analysis showed that 64 of these membrane proteins are localized on the cell surface and hence may serve as potential substrates for ADAM9. These include 35 single-pass transmembrane (29 type I, 3 type II, 1 type IV and 2 unknown), 26 multi-pass transmembrane, and 2 glycosylphosphatidylinositol-anchored proteins (Table S7).

### ALCAM, IL10RB and IGSF8 are ADAM9 substrates

We next selected several of the candidate substrates for validation. These included ALCAM, IGSF8, and IL10RB. KD of ADAM9 downregulated the shed ALCAM in the conditioned media and concomitantly upregulated the full-length proteins in the cell lysates (Fig. 3A). Conversely, overexpression of wild-type ADAM9, but not the protease-dead E348A mutant, enhanced shed ALCAM levels in the conditioned media (Fig. 3B), suggesting that ADAM9 protease activity is required for ALCAM shedding. Since ALCAM is known as a substrate for ADAM17 (23), we further examined if ADAM9 regulates ALCAM shedding indirectly via ADAM17 by using HCT116 cells with ADAM17 knocked out (Fig. 3C). In the complete absence of ADAM17, ectopic wild-type ADAM9 still enhanced the shedding of ALCAM, whereas the E348A mutant had markedly reduced activity (Fig. 3D), suggesting that ADAM9-mediated shedding of ALCAM is independent of ADAM17. In HCT116 cells with ADAM9 KD, we also observed inhibited shedding of the endogenous IGSF8, as shown by the decrease of shed IGSF8 ectodomain in the conditioned media and increase of full-length protein in the cell lysates (Fig. 4A). However, both wild-type ADAM9 and the E348A mutant appeared to promote IGSF8 shedding (Fig. 4B). Thus, IGSF8 is likely an indirect substrate for ADAM9. Although we were unable to detect the shed IL10RB in the conditioned media, the full-length protein was upregulated upon ADAM9 KD (Fig. 4C). Interestingly, the shed IL10RB ectodomain was found in the cell lysates when wild-type ADAM9, but not the E/A mutant, was overexpressed (Fig. 4D), implying that the ectodomain of IL10RB was retained on the cell-surface or internalized after shedding.

**Figure 3.**
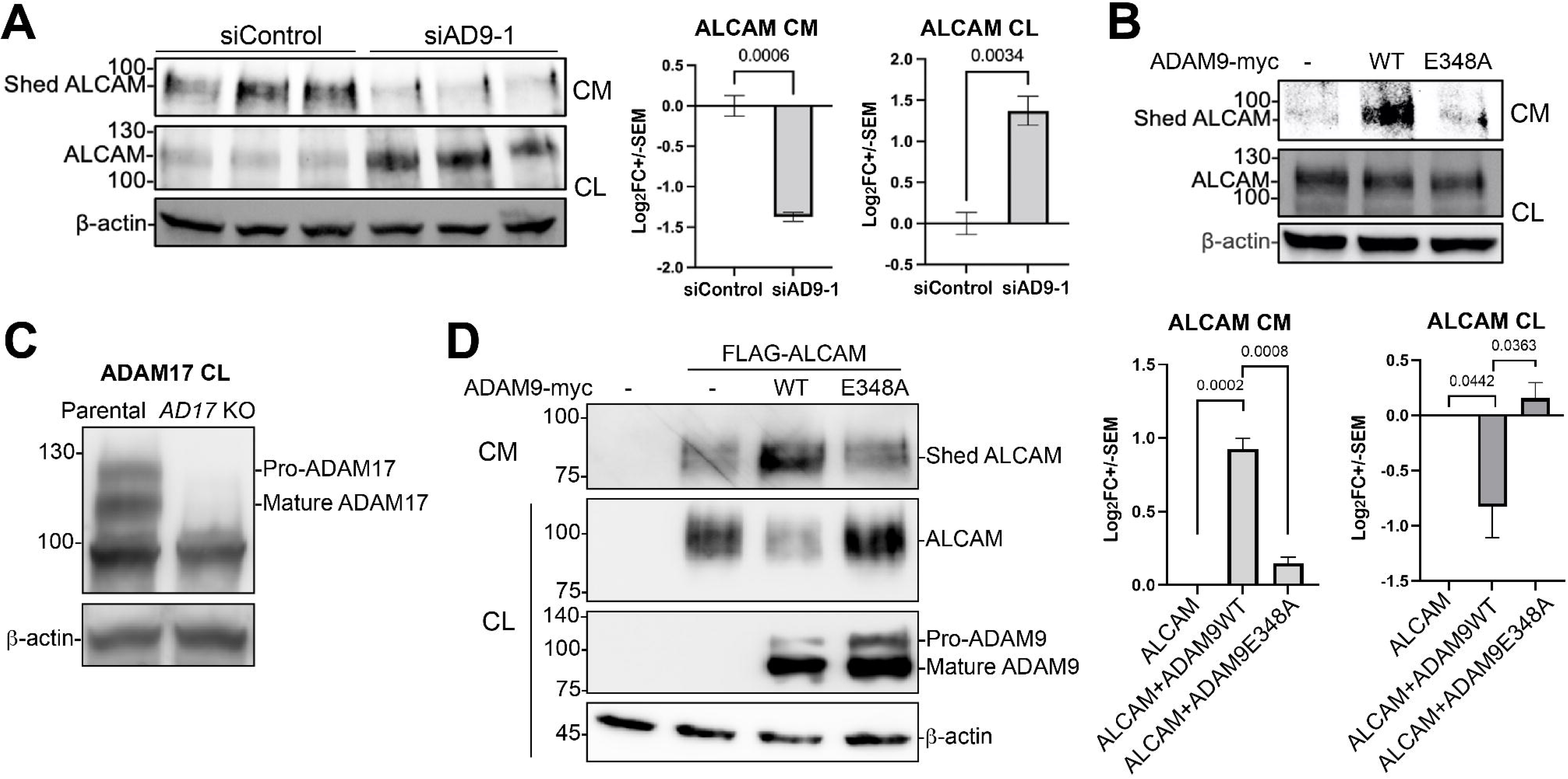
ADAM9 sheds ALCAM independently of ADAM17. (**A**, **B**) HCT116 cells were transfected with the indicated siRNA or expression constructs, and western blotting for 3 biological replicates was performed for conditioned media (CM) or cell lysates (CL) using an antibody recognizing the ectodomain of ALCAM. **(C)** Western blot of endogenous ADAM17 in parental and *ADAM17* KO HCT116 cells. **(D)** Shedding of extracellularly FLAG-tagged ALCAM by myc-tagged wild-type (WT) ADAM9 and the E348A mutant in *ADAM17* KO cells. Western blotting was carried out with an anti-FLAG antibody for ALCAM and an anti-myc antibody for ADAM9. Representative blots are shown on the left, and statistics including p-values on the right. Unpaired two-tail t test was performed for three biological replicates, and p-values are shown in **(A)** and **(D)**.

**Figure 4.**
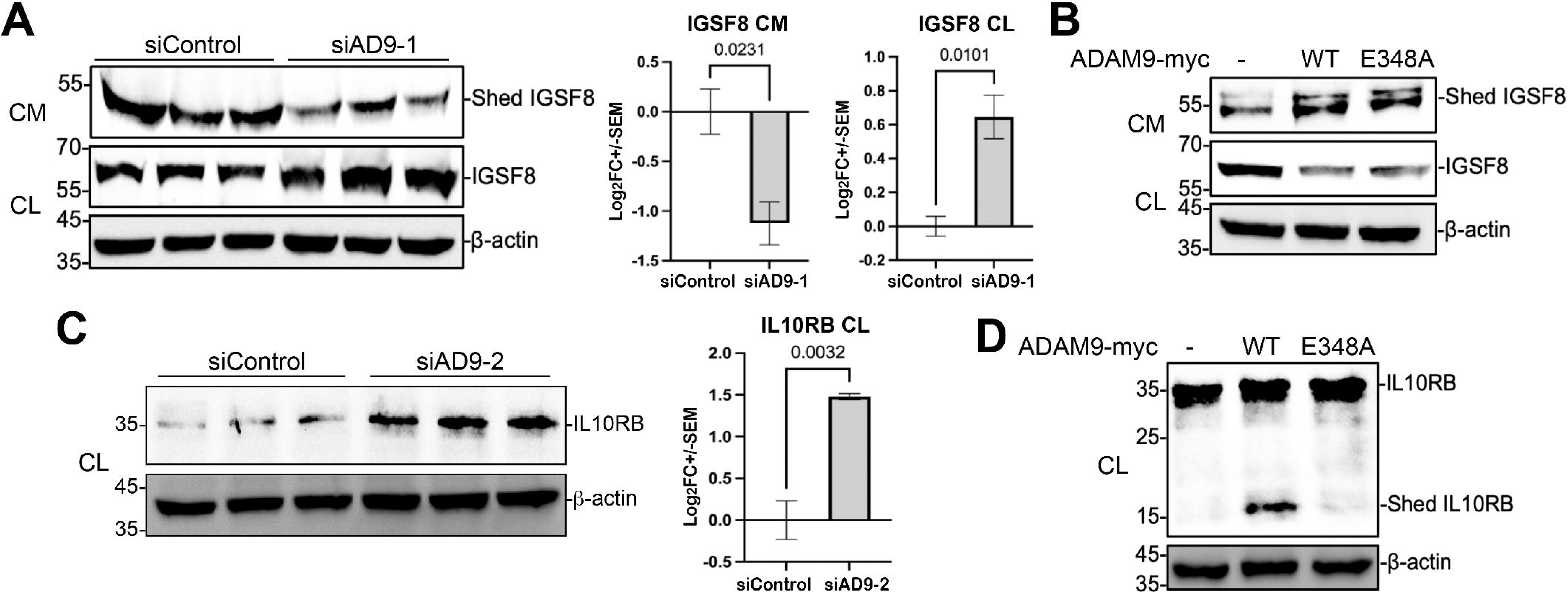
Identification of IGSF8 and IL10RB as new ADAM9 substrates. HCT116 cells were transfected with the indicated siRNA or expression constructs, and western blotting was performed for conditioned media (CM) or cell lysates (CL) using antibodies recognizing the ectodomain of IGSF8 **(A, B)** or IL10RB **(C, D)**. Three biological replicates are shown in **(A, C)**; unpaired two-tail t test was performed.

## Discussion

Given the importance of ADAM9 in various diseases, there is a critical need to identify its substrate repertoire and downstream targets. To date, only a handful of substrates have been reported for ADAM9, all of which were identified through candidate but not systematic approaches (10). Here we developed a new high-throughput and nonbiased approach to systematically screen for ADAM9 substrates. Among the validated substrates identified in this study, ALCAM and IGSF8 play important roles in solid tumor progression (24, 25). In particular, elevated ALCAM shedding in CRC samples correlates with poor patient survival (26). IL10RB is upregulated in both COPD and COVID-19 patients and is one of the top risk factors for severe COVID-19; it is also a potential therapeutic target for autoimmune diseases such as inflammatory bowel disease, which can lead to CRC (27–29). Of note, the shed IL10RB ectodomain was detected in the cell lysates but not the conditioned media, and cannot be identified using the traditional secretomics approaches. Since IL10RB primarily exists in complex with other receptor subunits such as IL10RA (30, 31), it is likely that IL10RB ectodomain shed by ADAM9 remains associated with these proteins on the membrane. We further show that ADAM9 promotes mTOR signaling, an oncogenic pathway (32), while suppressing FOXO signaling, a tumor suppressor pathway (33), providing an explanation for the involvement of this sheddase in tumor progression. It will be of interest to test if these substrates and targets mediate ADAM9 function in CRC and other related diseases.

Previously, secretomics and transcriptomics have been used as individual approaches to identify sheddase substrates and transcriptional targets, respectively (34–39). To our knowledge the integration of two or more omics approaches has not been applied to the studies of shedding. Here, we demonstrate the power of combining proteomics and transcriptomics to identify sheddase substrates and targets. Using ADAM9 as an example, we show that this method can differentiate between transcriptional and post-transcriptional targets, and reveal substrates that are not detectable by secretomics (such as IL10RB). Our future efforts will be focused on comparing and combining these results with those obtained from secretomics. The integration of multiple omics methods will greatly improve the coverage and accuracy of sheddase substrate and target identification.

## Experimental Procedures

### Plasmids and reagents

The expression constructs encoding wild-type and protease-dead E348A mutant ADAM9 were generated in our previous study (17). The cDNA construct for mouse ALCAM was purchased from Dharmacon (Clone ID: 3501215). The cDNA sequence encoding mouse ALCAM^28-583^ was subcloned into a pCS2+ expression vector containing a signal peptide followed by an in-frame FLAG tag using the BspEI and XbaI restriction sites (Fig. S5). Sequences of all PCR primers used in this study are shown in Table S8. The siRNAs used in this study include control siRNA (SIC) (Cell Signaling Technology (CST) 6568), ADAM9 siRNA-1 (AD9-1) (CST 11968), and ADAM9 siRNA-2 (AD9-2) (Ambion 4390826). Antibodies used in this study include rabbit anti-ADAM9 (CST 4151, 1:1,000), mouse anti-ALCAM (Santa Cruz 74558, 1:200), mouse anti-IL10RB (R&D Systems MAB874, 5 μg/ml), rabbit anti-FOXO3 (CST 2497, 1:1,000 for western blotting, and 1:100 For immunocytochemistry), rabbit anti-IGSF8 (Sigma 011917, 1:500), HRP-conjugated rabbit anti-mouse (CST 7076, 1:2,000), HRP-conjugated goat anti-rabbit (CST 7074; 1:2,000), HRP-conjugated mouse anti-β-actin antibody (CST 12262, 1:2,000), and Alexa Fluor™ 594-conjugated goat anti-rabbit antibody (Thermal fisher A-11012, 1:1,000 for ICC).

### Cell culture and transfection

Parental HCT116 were purchased from ATCC (CCL-247), and HCT116 cells with ADAM17 knocked out were from Abcam (ab266873). These cells were cultured in McCoy’s 5A medium (ATCC) containing 10% fetal bovine serum (FBS) and 1% Pen-Strep (100 U/mL penicillin and 100 μg/mL streptomycin) in 5% CO_2_ at 37°C. Cells were grown to 60% confluence and transfected with siRNA using lipofectamine RNAiMAX transfection reagent (Invitrogen). The medium was refreshed 12 h after transfection, and the cells were further incubated for another 48 h.

### Experimental design and statistical rationale

Six biological replicates of both control and ADAM9 KD cells were used for RNA-seq, and five biological replicates of both control and ADAM9 KD cells were used for proteomics, as recommended by published statistical studies (40, 41).

### RNA-seq analysis

Total RNA was extracted from the cells using RNeasy Mini Kit (QIAGEN, USA) according to the manufacturer’s instructions. The quality of the RNA was estimated using the A260:A280 ratio, and RNA degradation and contamination were monitored by electrophoresis in 1% agarose gels. The integrity and size distribution of RNA were evaluated using a Bioanalyzer. The total RNA samples were treated with DNase I to eliminate DNA contamination. Illumina RNA-Seq library construction and sequencing were conducted by the University of Delaware DNA Sequencing & Genotyping Center (RRID:SCR_012230, Newark, DE). Six replicates of each treatment were used for the RNA-seq (paried end with 101 read length) performed with the Illumina HiSeq 2500 system (Illumina).

After obtaining the RNA-Seq raw data in FASTQ format, the quality of the raw data was evaluated with FastQC (0.11.5) (Babraham Institute Bioinformatics, Cambridge, UK), a quality control tool for high throughput sequence data. The low-quality read ends and adapter sequences were removed, and the low-quality reads were filtered out using Trim Galore (0.4.4) (Babraham Institute Bioinformatics, Cambridge, UK). After the trimming and filtering, all the reads were aligned to the hg19 human reference genome using TopHat2 (2.1.1). HTseq (0.11) was used to count how many reads map to each annotated gene. The differential gene expression analysis was conducted using the R package edgeR (3.20.9), which provide statistical routines for determining differential expression based on a negative binomial model. False discovery rate (FDR) was used for multiple test correction, and the exact test in edgeR was conducted to determine the differentially expressed genes (DEGs) with a cutoff threshold of FDR < 0.05 and fold change > 1.5. Pathway enrichment analysis of the DEGs was performed using Enrichr.

### Cellular Proteomics Analysis

Proteins were reduced, alkylated, and purified by chloroform/methanol extraction prior to digestion with sequencing grade modified porcine trypsin (Promega). Tryptic peptides were labeled using tandem mass tag isobaric labeling reagents (Thermo) following the manufacturer’s instructions and combined into one 10-plex sample group. Labeling efficiency was >99%. The labeled peptide multiplex was separated into 46 fractions on a 100 x 1.0 mm Acquity BEH C18 column (Waters) using an UltiMate 3000 UHPLC system (Thermo) with a 50 min gradient from 99:1 to 60:40 buffer A: B ratio under basic pH conditions, and then consolidated into 18 super-fractions. Each super-fraction was then further separated by reverse phase XSelect CSH C18 2.5 um resin (Waters) on an in-line 150 x 0.075 mm column using an UltiMate 3000 RSLCnano system (Thermo). Peptides were eluted using a 60 min gradient from 98:2 to 60:40 buffer A: B ratio. Eluted peptides were ionized by electrospray (2.2 kV) followed by mass spectrometric analysis on an Orbitrap Eclipse Tribrid mass spectrometer (Thermo) using MultiNotch MS3 parameters. MS data were acquired using the FTMS analyzer in top-speed profile mode at a resolution of 120,000 over a range of 375 to 1500 m/z. Following CID activation with normalized collision energy of 35.0, MS/MS data were acquired using the ion trap analyzer in centroid mode and normal mass range. Using synchronous precursor selection, up to 10 MS/MS precursors were selected for HCD activation with normalized collision energy of 65.0, followed by the acquisition of MS3 reporter ion data using the FTMS analyzer in profile mode at a resolution of 50,000 over a range of 100-500 m/z. Buffer A = 0.1% formic acid, 0.5% acetonitrile, Buffer B = 0.1% formic acid, 99.9% acetonitrile. Both buffers were adjusted to pH 10 with ammonium hydroxide for offline separation.

Proteins were identified and MS3 reporter ions were quantified using MaxQuant (1.6.6.0) against the UniprotKB *Homo sapiens* database (January 2021; containing 20,394 protein entries) with a parent ion tolerance of 3 ppm, a fragment ion tolerance of 0.5 Da, and a reporter ion tolerance of 0.003 Da. Scaffold Q+S (Proteome Software) was used to verify MS/MS-based peptide and protein identifications (protein identifications were accepted if they could be established with less than 1.0% false discovery and contained at least two identified peptides; protein probabilities were assigned by the Protein Prophet algorithm) and to perform reporter ion-based statistical analysis (42). The scaffold output filtered the protein IDs to only those that have 2 or more peptides. Maximum missed cleavages permitted were set at 2. Fixed modifications were specified as carbamidomethylation (cysteine), and variable modifications were oxidation of methionine and protein N-terminal acetylation. Protein TMT MS3 reporter ion intensity values were analyzed using the ProteiNorm app, a user-friendly tool for normalization, imputation of missing values, and differential expression analysis (43). The Linear Models for Microarray Data (limma) with empirical Bayes (eBayes) smoothing to the standard errors were used to conduct the differential expression analysis (44). Proteins with an FDR < 0.05 and a fold change > 1.2 were considered to be differentially expressed proteins.

### KEGG pathway enrichment analyses

DEGs were analyzed for pathway enrichment using the Enrichr web tool (https://maayanlab.cloud/Enrichr/)(45, 46). A total of 351 DEGs, including both up- and down-regulated genes, were input into Enrichr, and enrichment analysis was performed using the KEGG 2021 human database under the “Pathways” category. Enrichr calculates enrichment using Fisher’s exact test and applies a background gene set consisting of all annotated human genes in the KEGG 2021 library. Pathways with raw p-values < 0.05 were considered significantly enriched.

### Analyses of overlap between two groups of genes

The statistical significance of the overlap between two groups of genes was calculated using http://nemates.org/MA/progs/overlap_stats.html. The number of protein-coding genes in the human genome (20,000) was used as the input for “number of genes in the genome”.

### Integration of transcriptome and proteome data to identify ADAM9 substrates and targets

The transcriptome and cellular proteome were integrated using the OmicsPLS R package (PMID: 30309317). OmicsPLS is the implementation of O2PLS, which is a symmetrical multivariate regression method that integrates two datasets (47, 48). It decomposes the variation in two data matrices into three parts: the joint component shared by two datasets, the orthogonal components specific for each dataset, and residual error. The join components induce the relationship between the two datasets and can be viewed as the representation of the integration. We define transcriptional targets as DEPs whose mRNA and protein positively correlate with each other, and post-transcriptional targets as DEPs whose mRNA and protein do not have a significant positive correlation (Fig. 1B). Pre-processed data was fitted to the O2PLS model to obtain the systematic and non-systematic components of the data, with the former including the joint component, and dataset-specific ‘orthogonal’ component. For both types of targets, we further determined whether they were positively or negatively correlated with ADAM9. To do this, we tested for a significant positive or negative correlation of the systematic variation of the protein with that of ADAM9 in the proteome. Potential substrates of ADAM9 cleavage are cell-surface proteins that are post-transcriptional targets negatively correlated with ADAM9. Cell Surface proteins were determined from their annotation as per UniProt (Release 2024_03) subcellular localization. The surface proteins were further prioritized based on their types.

### Western Blotting and RT-qPCR

Cells were lysed in the cell lysis buffer (Cell Signaling) and the concentration of the protein was determined using the Bicinchoninic acid (BCA) assay (Thermo Scientific) with bovine serum albumin (BSA) as standard. For the western blotting, the same amount of protein was loaded on SDS-PAGE gels and then electrotransferred onto PVDF membranes (Bio-Rad). After blocking for 1 hour with 5% BSA and incubating for 1 hour with primary antibody, signals were detected by peroxidase-conjugated secondary antibodies using chemiluminescence reaction (Bio-Rad). β-actin was used as the loading control.

Total RNA was reverse transcribed into cDNA using the iScript cDNA Synthesis Kit (Bio-Rad Laboratories) with DNase I treatment (Qiagen). RT-qPCR was performed using the qMAX Green Low Rox qPCR Mix (Accuris) according to the manufacturer’s instructions. All samples were assayed in triplicate, and data were analyzed using the Comparative CT Method with GAPDH as the reference gene. The RT-qPCR primers are shown in Table S8.

### Immunocytochemistry

HCT116 cells were grown to 60% confluence on Poly-L-lysine coated coverslips (Neuvitro) and transfected with siRNA using lipofectamine RNAiMAX transfection reagent (Invitrogen). The medium was refreshed 12 h after transfection, and the cells were further incubated for another 48 h. Cells were then fixed in PBS containing 4% paraformaldehyde for 10 min and permeabilized by incubation in PBS containing 0.2% Triton X-100 for 10 min at room temperature. After washing in PBS three times for 5 min, cells were blocked with 1% BSA for 30 min. Cells were then incubated with FOXO3 antibody (1:100) overnight at 4, washed in PBS three times for 5 min, and incubated with anti-rabbit secondary antibody (1:1000) for 1 h at room temperature. After washing in PBS three times for 5 min, coverslips were incubated in DAPI (1:10,000) for 1 min at room temperature. Imaging was performed with confocal microscopy LSM 880, AxioObserver, and objective Plan-Apochromat 20x. Nuclear regions were first defined by DAPI stain and the red signal intensities within the nuclear regions were measured.

## Data availability

Data supporting the findings of this work are available within the paper and its Extended View figures and tables files. The raw mass spectrometric data have been deposited at http://www.ebi.ac.uk/pride. Project accession: PXD059109; token: HMoBUNviKfsa. Or username; password: zJgLcf3DVFLI. RNA-seq data will be deposited to public repositories upon publication.

## Acknowledgements

This work was supported by the US National Institute of Health (R01 DE029802, P20 GM104316, U54 GM104941 and R24 GM137786 to S.W.). JR was supported by an NIH training grant (T32 GM133395).

