## Supplemental figures for "An integrative proteotranscriptomics approach reveals new ADAM9 substrates and downstream pathways"

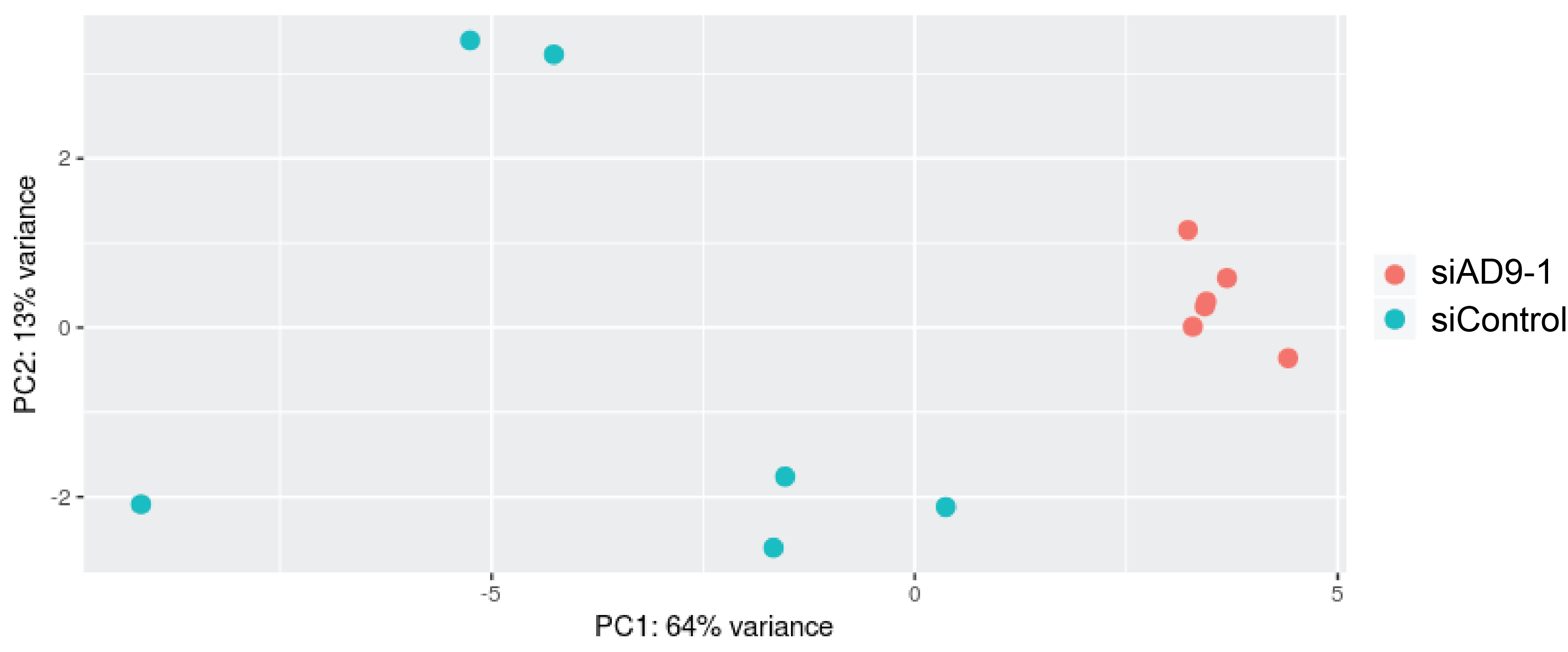

**Fig. S1. Principle component analysis (PCA) plot of RNA-seq data obtained from HCT116 cells treated with control or AD9-1 siRNA.**

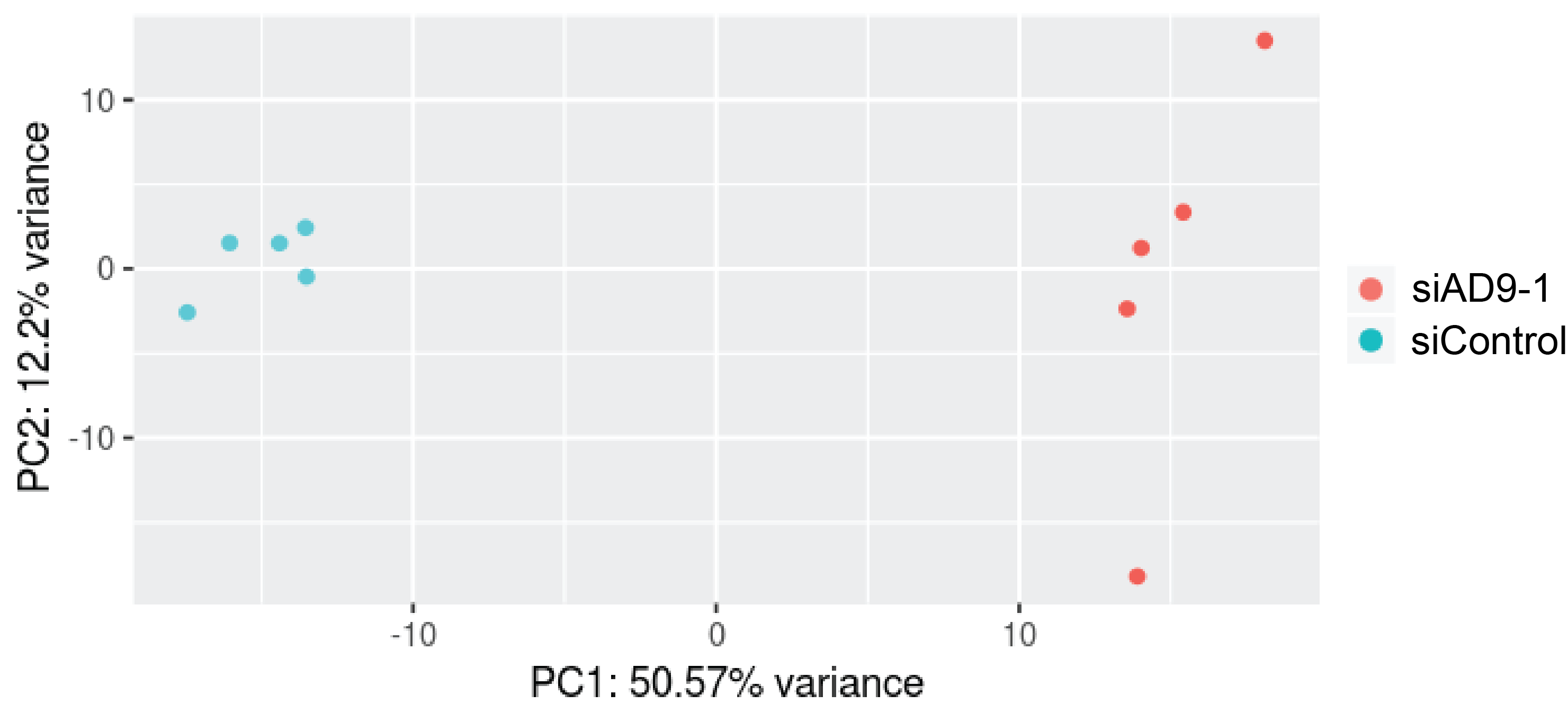

**Fig. S2. PCA plot of quantitative cellular proteomics obtained from HCT116 cells treated with control or AD9-1 siRNA.**

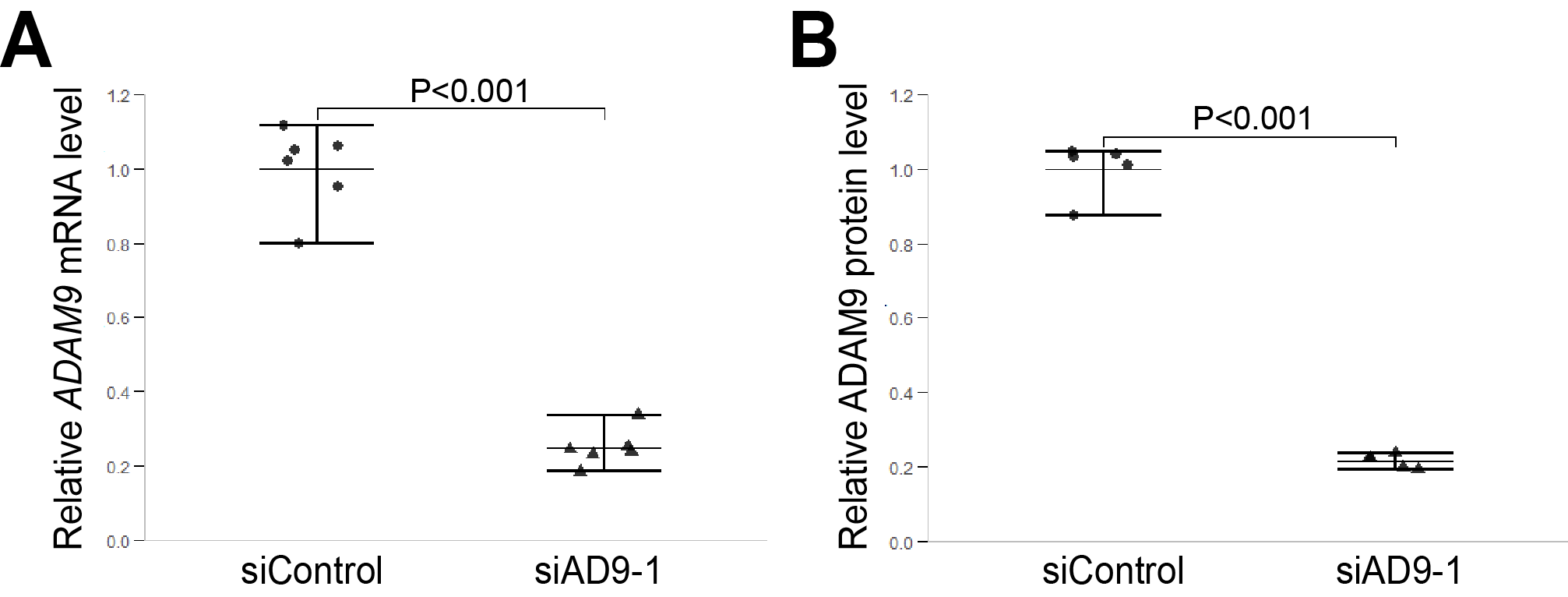

**Fig. S3. ADAM9 mRNA and protein levels in HCT116 cells treated with control or AD9-1 siRNA.** **A.** Relative *ADAM9* mRNA levels in RNA-seq samples (6 biological replicates each). **B.** Relative ADAM9 protein levels in proteomics samples (5 biological replicates each).

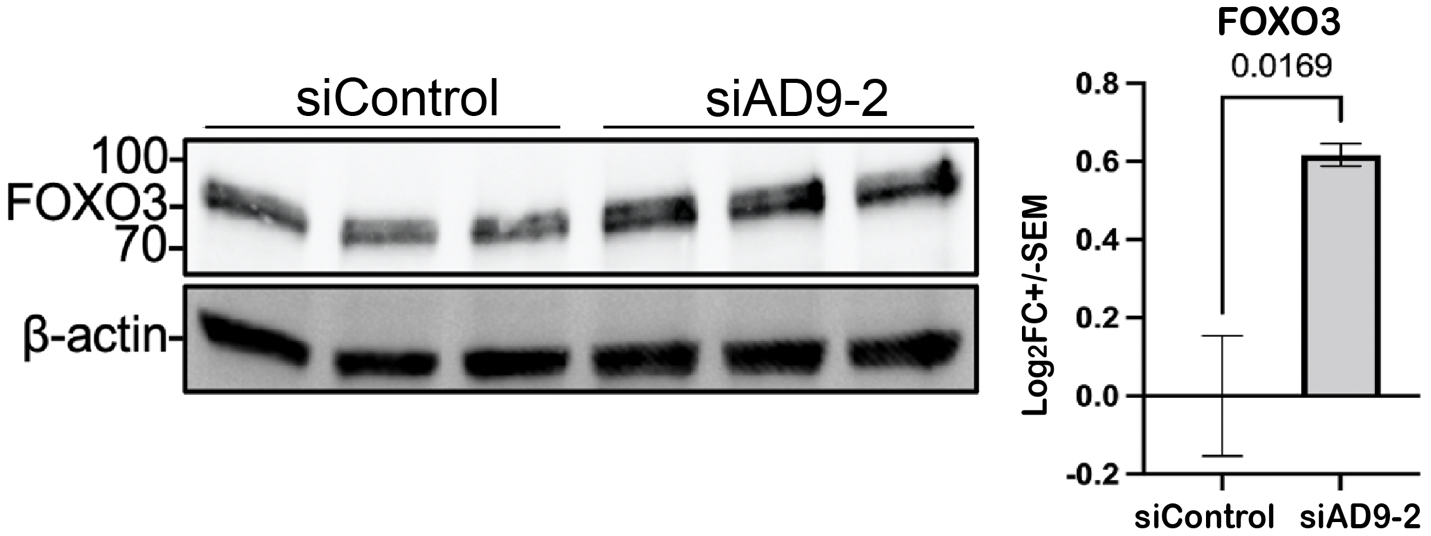

**Fig. S4. Western blots of FOXO3 in HCT116 cells treated with control or AD9-2 siRNA.** Cells were treated with the indicated siRNA (3 biological replicates each), and cell lysates were processed for western blotting with the indicated antibodies. Statistics is shown on the right with p-value at the top.

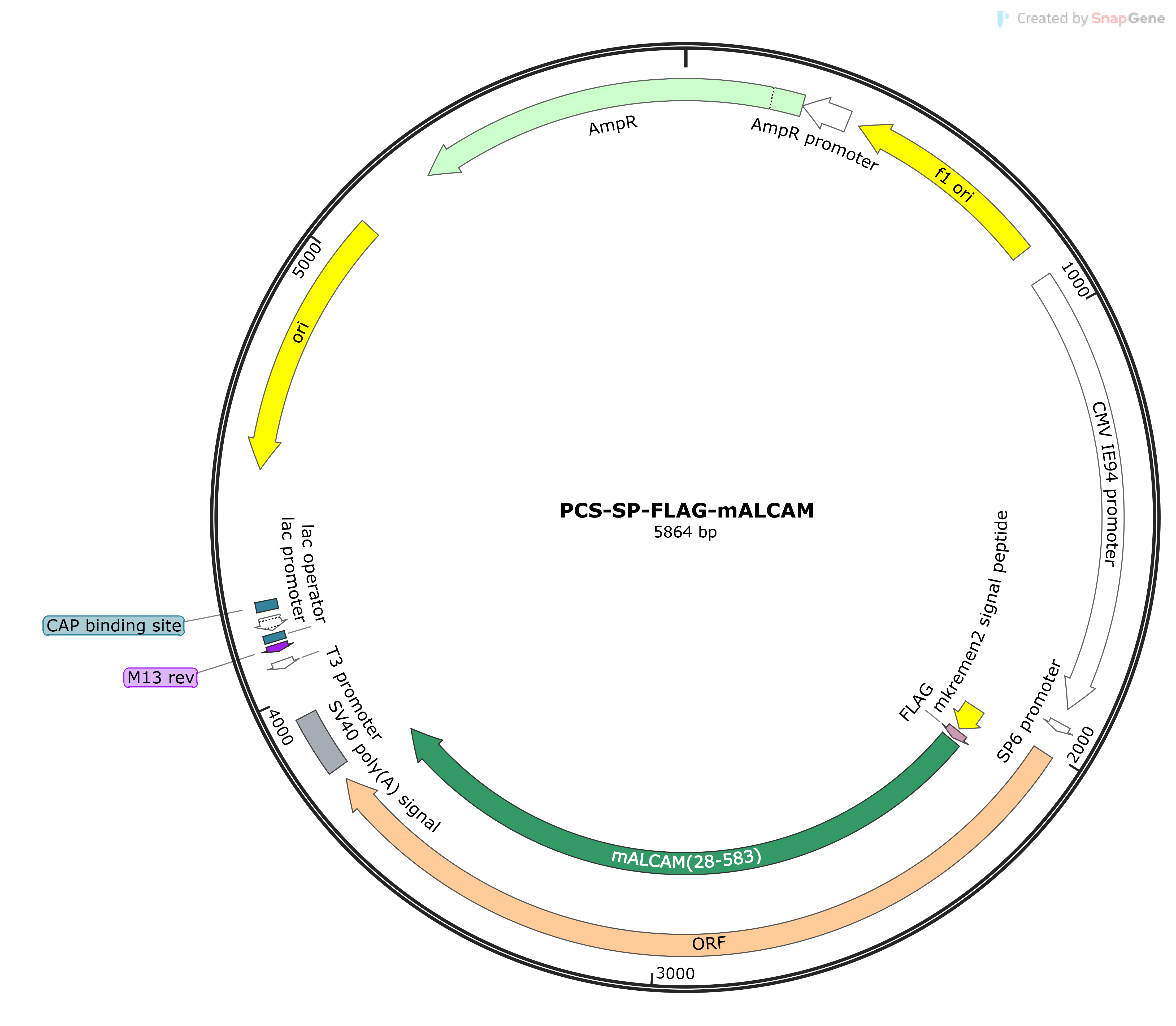

**Fig. S5. Map of the expression construct encoding N-terminally FLAG-tagged mouse ALCAM.**
